# Modulation of translational elongation by montanine alkaloids preferentially inhibits RNA viruses

**DOI:** 10.64898/2025.12.19.695327

**Authors:** Ryuichi Sakai, Kyohei Sato, Atsushi Tsugita, Lakkana Thaveepornkul, Ayato Takada, Hiroko Miyamoto, Rumi Kurokawa, Minoru Yoshida, Takeshi Yokoyama, Antonio Evidente, Yuichiro Tsuge, Hiromi Watari, Tomonosuke Sumiya, Ken Matsumoto, Sarin Chimnaronk, Yoshikazu Tanaka

**Affiliations:** Hokkaido University Faculty and Graduate School of Fisheries Sciences.; Graduate School of Life Sciences, Tohoku University, Sendai, Miyagi 980-8577, Japan; Institute of Molecular Biosciences, Mahidol University, 25/25 Phutthamonthon 4 Road, Salaya, Nakhon Pathom 73170, Thailand; Division of Global Epidemiology, International Institute for Zoonosis Control, Hokkaido University, Kita 20 Nishi 10, Sapporo 001-0020, Japan; Chemical Genomics Research Group, RIKEN Center for Sustainable Resource Science, 2-1 Hirosawa, Wako-Shi, 351-0198, Saitama, Japan; Drug Discovery Seeds Development Unit, Drug Discovery Platforms Cooperation Division, RIKEN Center for Sustainable Resource Science, 2-1 Hirosawa, Wako-Shi, Saitama 351-0198, Japan; Institute of Biomolecular Chemistry, National Research Council, Via Campi Flegrei 34, 80078 Pozzuoli (Na), Italy; Siriraj Center of Research Excellence for Systems Pharmacology, Department of Pharmacology, Faculty of Medicine Siriraj Hospital, Mahidol University, Bangkok 10700, Thailand

**Author notes:** **Author Contributions:** RS, YoT, and SC initiated the work. RS coordinated and supervised the study. SC, AyT, HM, and LT evaluated antiviral activities. KS, AtT, TY, and YoT performed the cryo-EM study, and KM, RK, and MY performed the chemical genomics experiments. ST, WH, and YuT conducted the cellular assays. RS, YT, SC, and KM analyzed the data. RS, YT, SC, and KM wrote the manuscript. AE isolated the natural alkaloids, isocarbostyryls and fungal metabolites, prepared the semis synthetic derivatives and edited the manuscript. **Competing Interest Statement:** Disclose any competing interests here. **Classification:** Paste the major and minor classification here. Dual classifications are permitted but cannot be within the same major classification.

## Abstract

Montanine, an Amaryllidaceae alkaloid, has demonstrated superior antiviral activity against dengue virus (DENV), SARS-CoV-2, vesicular stomatitis Indiana virus (VSV-G), and vesicular stomatitis virus pseudotyped with Ebola virus glycoprotein (VSV-ZGP), outperforming other antiviral alkaloids tested, including lycorine, narciclasine, tetracetylnarciclasine, and pancracine. We further showed that montanine inhibits translation in mammalian cells, as evidenced by a puromycin incorporation assay. Cryo-EM analysis revealed that montanine binds to the peptidyl transferase center of the human ribosome. A chemical genomics survey indicated that the knockdown of the GCN1 complex, which senses ribosome collisions and triggers the translation quality control process, increased the cytotoxicity of montanine while reducing viral infectivity. Together, these results suggest that the antiviral activity of montanine is closely linked to its impact on translational elongation and GCN1-related stress responses, underscoring the potential of translation control as a targeted mode for RNA virus therapeutics.

**Significance Statement:** Montanine, an alkaloid from the Amaryllidaceae family known for its diverse bioactivities, was identified as a potent antiviral compound from a library of natural plant- and fungus-derived substances. Through a combination of biochemical assays and cryo-EM analysis, we discovered that montanine binds to the peptidyl transfer center of the human ribosome, effectively inhibiting protein synthesis. Based on these findings and a chemical genomics approach, we hypothesize that the translational control of montanine triggers a cellular response that impedes viral replication. We propose that molecules capable of modulating translation efficacy could serve as unique lead compounds in the development of broad-spectrum antiviral agents.

## Introduction

Infectious diseases, especially viral infection, impact not only human health but also social and economic stability because of their sporadic nature, high ability for transmission and serious medical consequences. COVID-19, for example, first emerged in 2019 in Wuhan, China, but rapidly spread worldwide in several months, and to date, 775 million cumulative cases and 7 million deaths have been reported by the World Health Organization. The economic and health loss caused by this pandemic was estimated to be $16 trillion (1). The COVID-19 pandemic clearly illustrates the vulnerability of modern society to viral diseases and highlights the need to prepare against the next yet unknown pandemic. To this end, vaccine development would be a general and the most efficacious strategy if a pathogen virus, such as influenza, was previously identified. However, vaccine development might be difficult for some viruses. For example, vaccine development for dengue virus (DENV) has been hampered by the immunological complexity of the viruses where four immunologically different serotypes (DENV-1–4) exist; thus, a vaccine is needed to suppress infection with all four serotypes together to avoid antibody-mediated enhancement of infection, which may cause severe symptoms (2). Recently, a newly developed dengue vaccine, TAK-003, has shown promising efficacy in controlling DENV-1 and DENV-2, but its efficacy against DENV-3 and DENV-4 is still unclear, and further evaluation is necessary to characterize this vaccine (3). The development of vaccines for unpredictable but highly aggressive viruses such as viral hemorrhagic fever virus (VHF), including the Ebola virus, is rather challenging, as the occurrence of this virus is unpredictable, the number of patients is limited, and there is a lack of investment in the development of clinically useful and commercially feasible vaccines. Recently, however, several vaccines and neutralizing antibodies have been developed, including a viral vector-based vaccine, rVSV-ZEBOV (4), and two monoclonal antibodies, which have received FDA approval (5); however, evaluating their actual efficacy remains difficult (6, 7).

Small-molecule antiviral drugs that prevent infection or mitigate symptoms during early viral infection serve as the first-line defense in antiviral therapy. The recent discovery of the anti-DENV drug JNJ-1802 demonstrated the promising future of modern small-molecule strategies for highly challenging disease targets (8). Therefore, continuous efforts to develop small antiviral drugs have been proven to be important for increasing preparedness for future unexpectable pandemics (9). To this end, a broad-spectrum antiviral drug (BSA) strategy (10) rather than a virus-one drug approach (11) is preferable. Currently, a total of 150 compounds have been listed as BSAs, and knowledge-based accumulation of the structural entity that has broad-spectrum antiviral activity is available (12).

In the present study, we explored our natural product library to discover both virus-specific BSAs and BSAs using DENV, SARS-CoV-2, and vesicular stomatitis Indiana virus (VSV-G) and VSV pseudotyped with Ebola virus glycoprotein (VSV-ZGP) and found that several Amaryllidaceae alkaloids (AAs) and close isocarbostyrils (ICs) are candidates for novel BSAs. Amaryllidaceae plants are known to be rich sources of bioactive alkaloids (13), and compounds of this class, represented by lycorine (**1**), are known to possess antiviral activities (14). Here, we describe the screening results that led to the discovery of montanine as a potent antiviral agent. We identified montanine as an inhibitor of translation in human cells and determined the structure of the human ribosome in complex with montanine at 2.8 Å resolution via cryo-electron microscopy (cryo-EM). We further discuss the possible mode of antiviral action of montanine on the basis of the results from chemical genomics experiments.

## Results

### Antiviral activities of plant- and fungus-derived natural products: A primary screening

We first screened for antiviral activities on our pure compound library derived from terrestrial plants and fungal metabolites to determine the relationships among chemical structure, molecular targets, and antiviral actions. The compounds tested in the present study are shown in Figure 1. The sources and previously known bioactivities of these compounds are summarized in Table S1. Compounds **5**-**13** and **16**-**28** were subjected to primary screening with the viruses including DENV, SARS-CoV-2 and VSV-ZGP. VSV-G, a vehicle of pseudotype virus was used as control.

**Figure 1.**
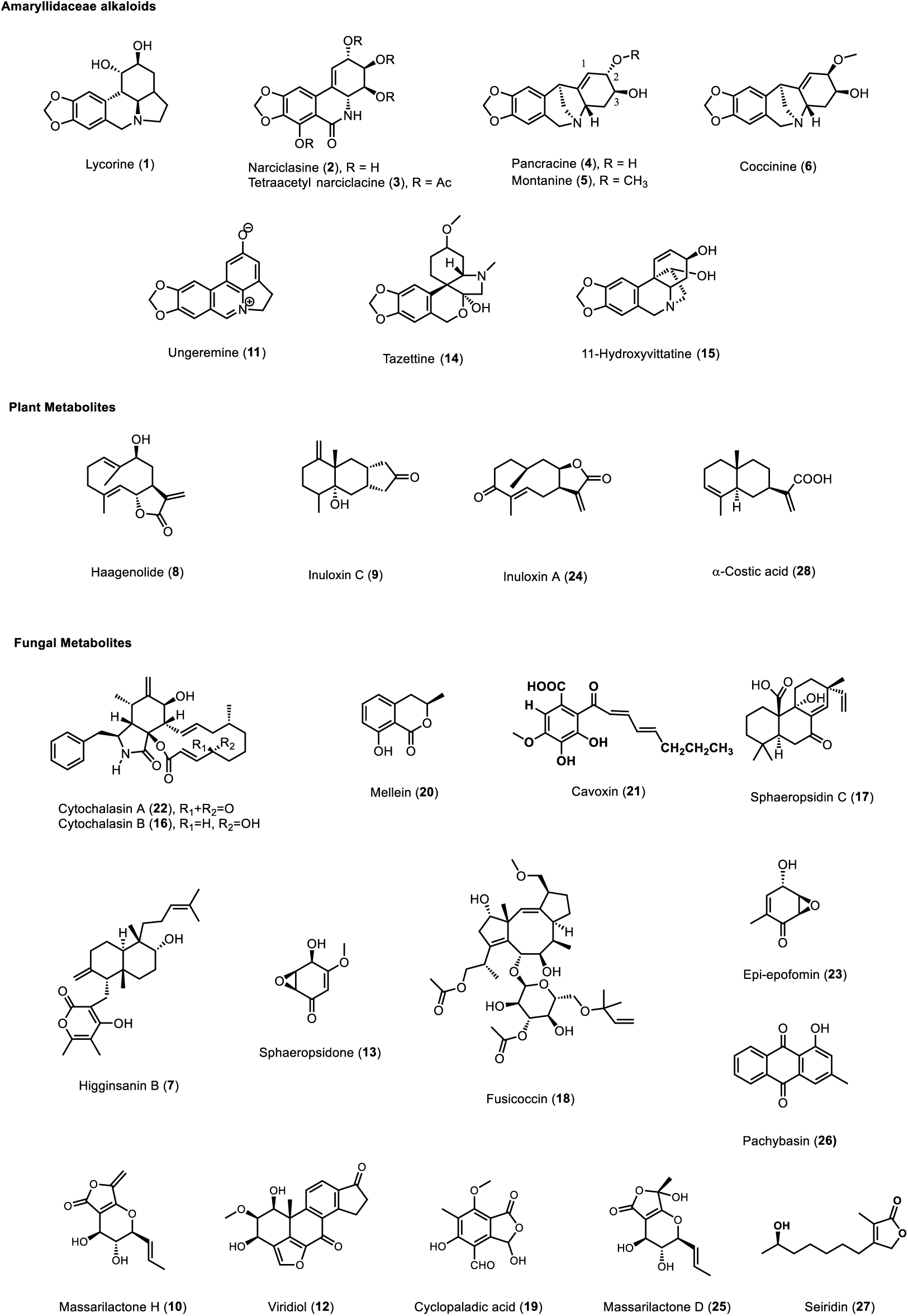
Chemical structures of natural products used in the present study.

First, anti-DENV activity was surveyed in HuH-7 cells infected with DENV-2. Each compound was applied to the cells (10 μg/mL), and viral RNA replication was evaluated via high-content analysis (HCA) with an anti-double-stranded RNA (dsRNA) antibody. Note that limited sample amount permitted the screening assays to test with n=1 but the rather qualitative result was satisfactory to select the candidates. Cell viability was assessed by DAPI nuclear staining. The activity profiles were mapped according to viral inhibition and cell viability and categorized into four groups (Figure 2A). Among the 22 compounds tested, two AAs, montanine (**5**) and coccinine (**6**) (15), showed nearly 100% inhibition, with more than 30% cell viability (Group A, Figure 2A). Fluorescent image clearly showed cell protection by **5** and **6** with these compounds (Figure 2B). Group B, which presented moderate viral inhibition (39–50%) and cell viability (9–30%), was composed of higginsanin B (**7**), haagenolide (**8**), inuloxin C (**9**) and massarilactone H (**10**). *N*-(4-hydroxyphenyl)retinamide (4-HPR or fenretinide), used as a positive control at 10 µM, fell into this category. Ungeremine (**11**) and viridiol (**12**), which displayed high viral inhibition but low cell viability, were categorized into group C. All the other compounds in group D presented negligible antiviral activity (Figure 2A).

**Figure 2.**
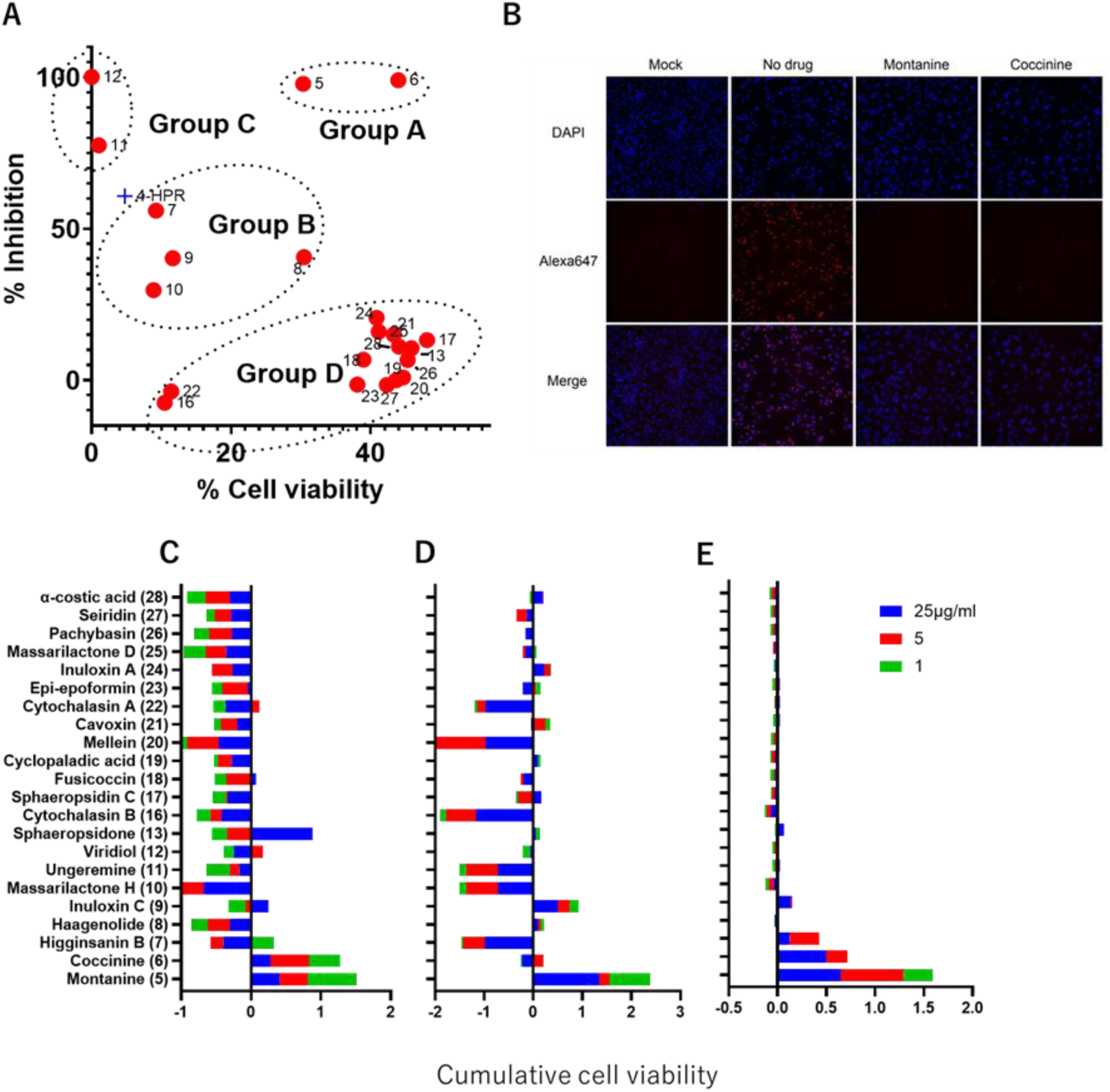
Antiviral screening of the first compound library (compounds 5–13, 16–28). (**A**) Inhibition of DENV-2 infection versus cell viability in HuH-7 cells. (**B**) Fluorescence micrographs of infected cells. Red and blue fluorescence indicate DENV-2 double-stranded RNA stained with an anti-dsRNA antibody and cell nuclei stained with DAPI, respectively. (**C**) Cumulative cell-viability values (sum of MTT assay data) against VSV-ZGP and SARS-CoV-2 in Vero cells at three concentrations (25, 5, and 1 µg/mL). Only compounds exhibiting more than 10% cell protection are shown.

We next evaluated the antiviral activity of the above compounds against VSV-ZGP, VSV-G, and SARS-CoV-2 (Figure 2C-E). In this assay, a test compound and virus were applied simultaneously to Vero E6 cells, and cell viability was assessed via the MTT assay at three concentrations (25, 5, and 1 μg/mL). The cytopathic effect (CPE), an indicator of cell damage caused by viral infection, was observed under a microscope and recorded individually. The compound that suppressed CPE was considered antiviral candidate. Figure 2C-E indicates cumulative cell viability values across all the concentrations tested for VSV-ZGP, SARS-CoV-2 and VSV, respectively. We found that **5** exhibited protective potential against virus-induced cell damage at all the concentrations tested indicated by reduction of CPE and retained cell growth. Coccinine (**6**) supported cell growth in VSV-ZGP with some reduction of CPE. In SARS-CoV-2 6 poorly supported cell growth although CPE reduction was observed at 5μ g/mL. In VSV **6** supported cell growth but no CPE reduction was observed across all concentrations tested. Generally all other compounds tested in the present study failed to protect cells from viral infection but rather toxic except for Sphaeropsidone (**13**) witch partially protected VSV-ZGP-treated cells at the highest concentration tested. Above results together demonstrated that montanine (**5**) and coccinine (**6**) are potential broad spectrum antiviral agents.

Montanine (**5**) and coccinine (**6**) both are AAs. Because some AAs are known to be potent antiviral agents, we tested the antiviral actions of **5** and **6** in detail along with six additional AAs and ICs with known antiviral activities, including lycorine (**1**), narciclasine (**2**), pancracine (**4**), tazettine (**14**) (16), 11-hydroxyvittatine (**15**) (17) and a derivative of **2**, tetraacetylnarciclasine (**3**), whose antiviral activity has yet to be reported (Table 1). All the compounds, except **2** (8.83 μM), did not show strong cytotoxicity against Vero E6 cells at the highest concentration tested. Lycorine (**1**), a well-known antiviral alkaloid, exhibited only moderate inhibitory activity against SARS-CoV-2 and the pseudotyped Ebola virus. Narciclasines **2** and **3** showed potent activity against both SARS-CoV-2 and VSV-ZGP, with IC_50_ values ranging from 71 to 381 nM. The montanine-class compounds **4**, **5**, and **6** also presented varying degrees of antiviral activity, of which **5** displayed robust inhibition of both viruses, with IC_50_ values of 1.71 and 0.277 μM, respectively. Our results indicated that **3** and **5** were more favorable broad antiviral agents because of their nanomolar IC_50_ values and higher selectivity index (SI) values beyond 90 (i.e., high efficacy with low cytotoxicity).

**Table 1.**
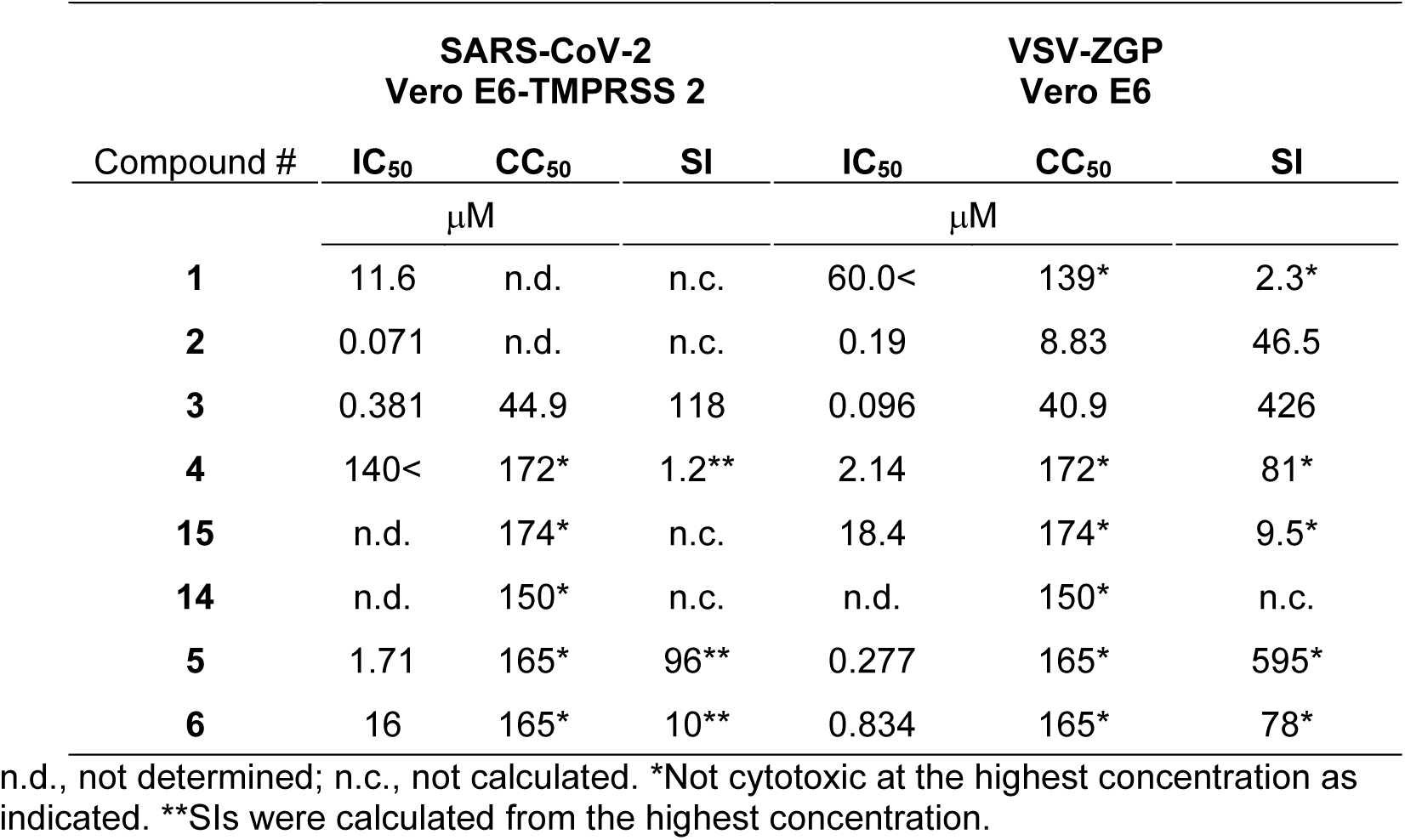
Antiviral activity, cytotoxicity, and selectivity indices of selected AAs and ICs.

| Compound # | SARS-CoV-2<br>Vero E6-TMPRSS 2 |  |  | VSV-ZGP<br>Vero E6 |  |  |
| --- | --- | --- | --- | --- | --- | --- |
|  | IC<sub>50</sub> | CC<sub>50</sub> | SI | IC<sub>50</sub> | CC<sub>50</sub> | SI |
|  | μM |  |  | μM |  |  |
| **1** | 11.6 | n.d. | n.c. | 60.0< | 139* | 2.3* |
| **2** | 0.071 | n.d. | n.c. | 0.19 | 8.83 | 46.5 |
| **3** | 0.381 | 44.9 | 118 | 0.096 | 40.9 | 426 |
| **4** | 140< | 172* | 1.2** | 2.14 | 172* | 81* |
| **15** | n.d. | 174* | n.c. | 18.4 | 174* | 9.5* |
| **14** | n.d. | 150* | n.c. | n.d. | 150* | n.c. |
| **5** | 1.71 | 165* | 96** | 0.277 | 165* | 595* |
| **6** | 16 | 165* | 10** | 0.834 | 165* | 78* |

n.d., not determined; n.c., not calculated. \*Not cytotoxic at the highest concentration as indicated. \*\*SIs were calculated from the highest concentration.

These results prompted further evaluation of the antiviral activity of the AAs against all four serotypes of DENV (DENV-1–4) in hepatic HuH-7 cells (Table 2). While all AAs tested clearly showed anti-DENV activity for all serotypes, some notable structure-activity relationship within the montanine-type alkaloids, **4**-**6** was observed, that is, **5** and **6**, its epimer at C2, showed provided the highest SI values across the 4 serotypes, while **4**, a desmethyl derivative of **5**, exhibited significantly weaker anti-DENV activity against DENV-1–4, suggesting that the methyl group in **5** was essential for anti-DENV activity. The results demonstrated that compounds **3** and **5** exhibited the highest potency, with submicromolar IC_50_ values against all DENV serotypes. Notably, montanine (**5**) emerged as the most promising broad-spectrum antiviral agent in this study, showing activity against SARS-CoV-2 and VSV-ZGP in addition to DENV.

**Table 2.** Anti-DENV activity of Amaryllidaceae alkaloids and isocarbostyrils.

| Compound # | DENV-1 |  | DENV-2 |  | DENV-3 |  | DENV-4 |  | HuH-7 |
| --- | --- | --- | --- | --- | --- | --- | --- | --- | --- |
|  | IC<sub>50</sub> | SI | IC<sub>50</sub> | SI | IC<sub>50</sub> | SI | IC<sub>50</sub> | SI | CC<sub>50</sub> |
|  | μM |  | μM |  | μM |  | μM |  | μM |
| **1** | 0.42 | 9.5 | 0.72 | 5.5 | 0.27 | 14.7 | 0.33 | 12.0 | 3.97 |
| **3** | 0.11 | 5.3 | 0.20 | 2.9 | 0.07 | 8.3 | 0.13 | 4.5 | 0.58 |
| **4** | 1.81 | 0.7 | 2.71 | 0.5 | 0.93 | 1.4 | 0.66 | 2.0 | 1.33 |
| **15** | 8.47 | 1.0 | 19.60 | 0.4 | 5.65 | 1.5 | 5.16 | 1.7 | 8.69 |
| **5** | 0.26 | 10.3 | 0.40 | 6.7 | 0.18 | 14.8 | 0.06 | 44.5 | 2.67 |
| **6** | 0.94 | 16.5 | 1.60 | 9.7 | 0.69 | 22.5 | 0.28 | 55.4 | 15.50 |

### Mechanism of antiviral activity

Given the broad antiviral activity of montanine-type compounds, they are likely to target the host machinery rather than viral proteins. We thus performed a LysoTracker assay (18) to detect acidification of endosomes to determine whether montanine interferes with viral entry by inhibiting endosomal acidification. However, we found that montanine had no effect on acidification, excluding its deacidification of the endosome lumen, as observed with entrance inhibitors (Figure S3). Because representative AAs and ICs, including lycorine (**1**) and narciclasine (**2**), are known to inhibit protein synthesis (19), we assessed protein synthesis inhibition by some AAs. We used the puromycin incorporation assay (20) adopted for ELISA. Because puromycin is incorporated into newly synthesized proteins by mimicking tRNA, measurement of its incorporation using an anti-puromycin antibody indicates translational activity. The results revealed that montanine also inhibited protein synthesis at 1 and 3 μM, but the inhibitory activities of coccinine and pancracine were not drastic or even negligible at 3 μM, respectively (Figure 3).

**Figure 3.**
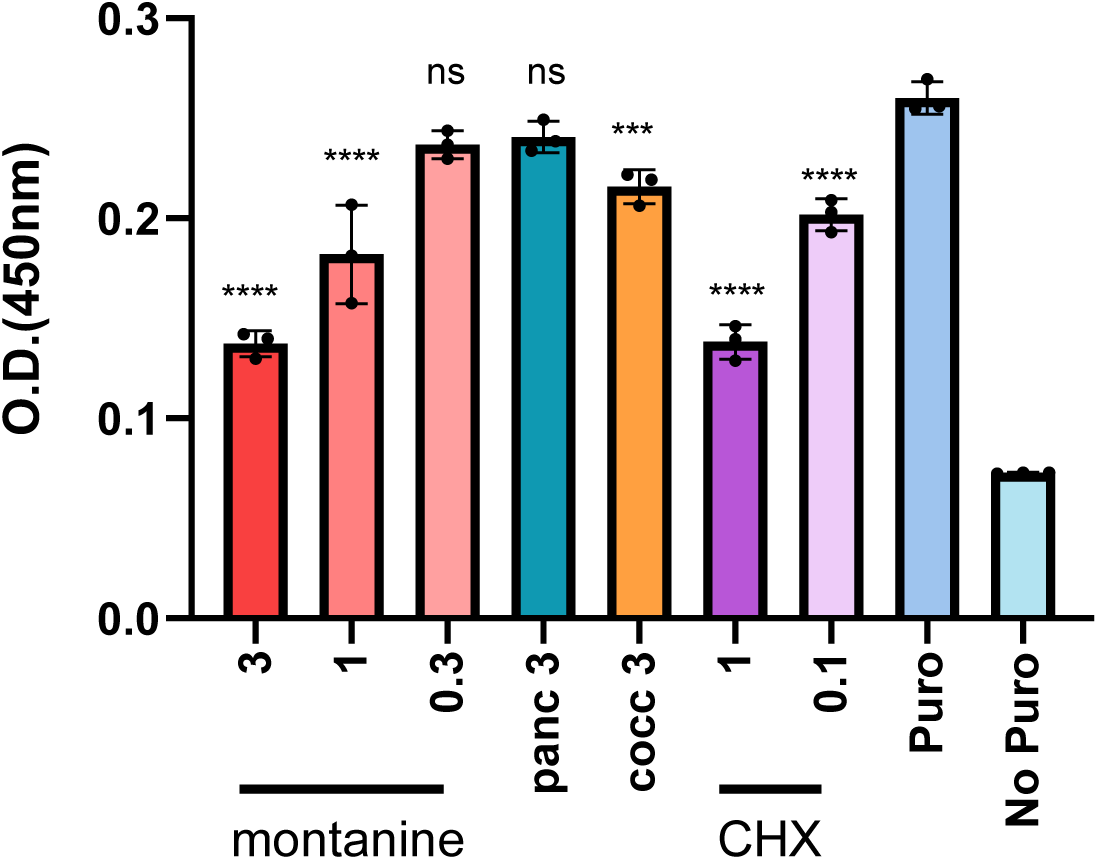
Inhibition of protein synthesis by montanine-type alkaloids was assessed by puromycin incorporation assay with an anti-puromycin antibody. Three and two concentrations (µM) of montanine and cycloheximide (CHX), respectively, and fixed concentrations (3 µM) of pancracine (panc) and coccinine (cocc) are shown. *P* (vs Puro): **** <0.0001, *** 0.0006, ns >0.05. Details of the statistical analysis performed via one-way ANOVA followed by Dunnett’s multiple comparison test are provided in Figure S4.

### Structural determination of montanine complexed with human ribosomes by cryo-EM single-particle analysis

It was previously reported that both **1** and **2** bind to the peptidyl transferase center (PTC) of the ribosome (21), prompting us to examine whether **5** has an affinity for the ribosome, although the structure of montanines differs largely from those ligands. We successfully determined the cryo-EM structure of the human ribosome in the presence of 30 µM montanine at an overall resolution of 2.8 Å. In the revealed structure, a clear density derived from montanine (**5**) was found at the PTC, in which the peptide attached to the 3’-end of the P-site tRNA was transferred to the amino acid attached to the A-site tRNA (Figure 4A). Montanine was located in a pocket composed of G3907, C4398, A4449, U4450, and U4452 and was accommodated via π‒π interactions and hydrogen bonds with the bases of U4398 and U4452 (Figure 4A and S5). A hydroxyl group at C3 of the cyclohexene ring (Figure 1) formed a hydrogen bond with O4 of U4452 (3.7 Å) as well as polar interactions with O6 (3.5 Å) and N1 (3.5 Å) of G3907. The ether oxygen of the methoxy group at C2 of the cyclohexene ring was in close proximity to the hydroxy group (O2’) of U4450 (3.3 Å) and the oxygen (O5’) involved in the phosphodiester bond of G4451 (3.5 Å). Moreover, the methoxymethyl group was also closely located to a hydroxy group (O2) of A4449 (3.5 Å) (Figure 4B). These atomic interactions within differentially polarized atoms (22) form a noncovalent network to stabilize the binding of montanine (**5**) to the pocket. The substantially lower activity of coccinine (**6**) and pancracine (**4**), attributed to the altered stereochemistry of the methoxy groups in the former and the substitution of a hydroxy group for a methoxy group at C-2 in the latter, may result in their inability to form these interactions. The binding site of montanine (**5**) coincided with the CCA terminus of the A-site tRNA, where amino acids are present during peptide transfer (Figure 4A, C); thus, the binding of montanine clashes with an incoming aminoacyl-tRNA (Figure 4D).

**Figure 4.**
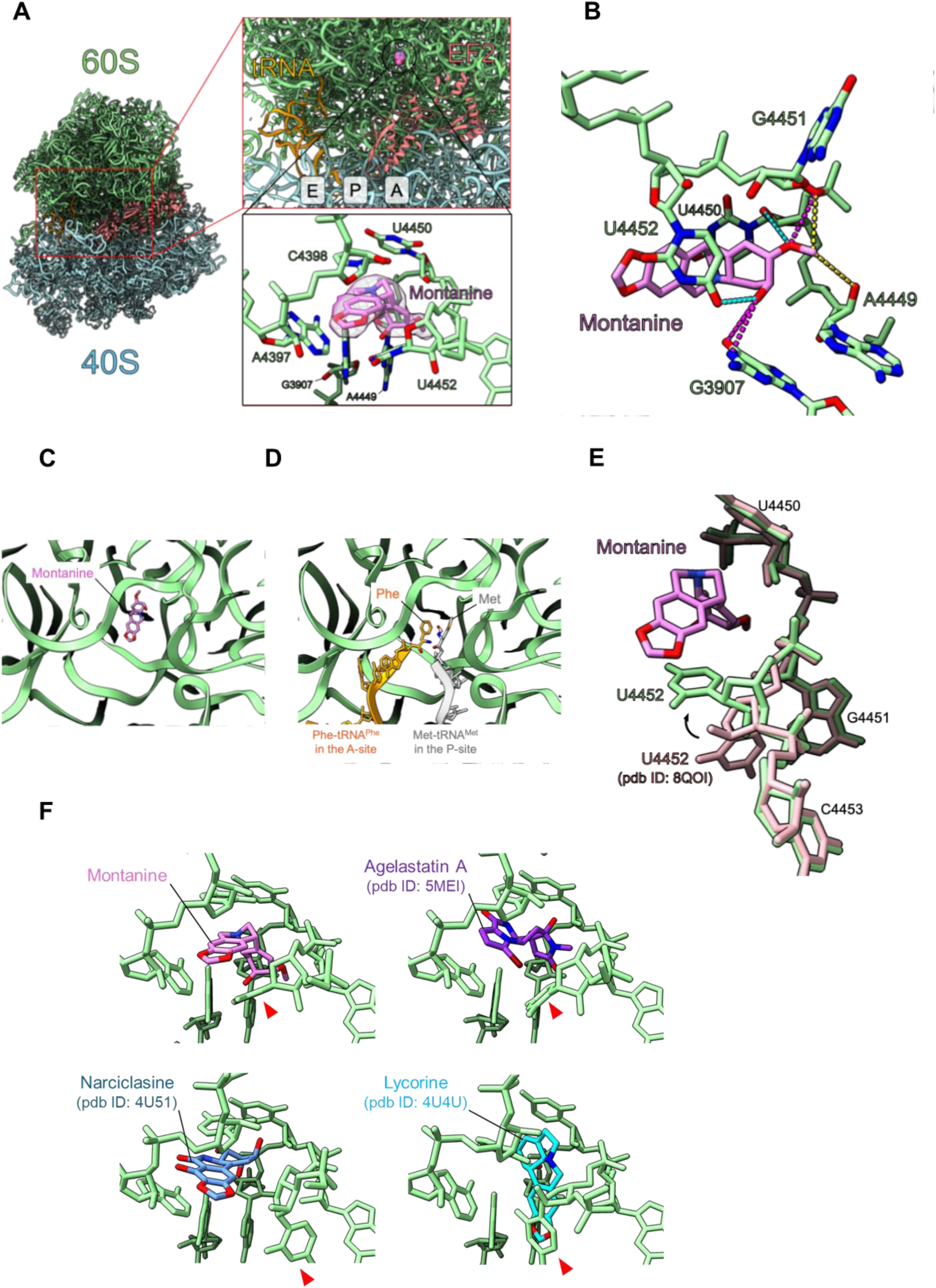
Cryo-EM analysis of the human 80S ribosome in complex with montanine. (**A**) Overall structure (left) and close-up images of the E-, P-, and A-sites (top right) and the montanine binding site (right bottom). The 60S subunit, 40 S subunit, tRNA, EF2, and montanine are shown in green, cyan, orange, salmon, and pink, respectively. Residues that interact with montanine are indicated. (**B**) Interactions involving the methoxy and hydroxy groups of montanine. Hydrogen bonds (blue), polar interactions (pink), and van der Waals interactions (yellow) are shown as dotted lines. (**C**) The relative locations of the montanine binding site and the amino acids at the CCA terminus of tRNAs. Montanine on the human 80S ribosome (pink) is shown. (**D**) Structure of the PTC immediately before transpeptidation (PDB ID 6WDD). tRNA^Phe^ at the A-site (yellow) and tRNA^Met^ at the P-site (gray) are shown. (**E**) Conformational changes in ribosomes upon montanine binding. pink: montanine, green: montanine complex, salmon: ligand-free human 80S ribosome (8QOI) (**F**). Close-up view of the PTC of the human 80S ribosome complexed with montanine and that of the yeast 80S ribosome complexed with agelastatin (5MEI), narciclasine (4U51), and lycorine (4U4U). Red arrows indicate U4552.

Binding with montanine (**5**) induced a large conformational change in U4452 (Figure 4E), likely due to the formation of sandwich-type π-π interactions and hydrogen bonds (Figure S5). The structural counterpart of U4452 in *E. coli* (U2506) plays an important role in peptide release and binding of P-site tRNA, and mutation of U2506 reduces transpeptidation activity (23). Several naturally occurring A-site inhibitors, including **1**, **2**, homoharringtonine, nagilactone C, and anisomycin, and a few mycotoxins, such as T-2 toxin (21), bind to the same site as montanine (**5**) does, even though the structures differ greatly from those of montanine (**5**) (Figure 4F). The marine antitumor alkaloid agelastatin A was also found to bind to the same site (Figure 4F) (24). Interestingly, a conformational change in U4452 upon binding of **5** was also found in the binding of agelastatin A and, to some extent, anisomycin (24), but no drastic change was observed in other inhibitors, including **1** or **2** (Figure 4E, F).

Taken together, these observations suggested that montanine likely inhibited peptidyl transfer at the PTC of the ribosome via steric competition with aminoacyl-tRNA at the A-site. The relationships between A-site inhibitors with different antiviral potencies and their binding modes to eukaryotic ribosomes are intriguing.

### Target identification by chemical genomics

It was not clear why montatine-mediated A-site inhibition could have a greater impact on viral replication than on host translation, as shown by the SI values in our antiviral assays. To identify the specific mechanism or pathways contributing to the potent antiviral action of montanine (**5**), we employed chemical genomic screening, in which approximately 15,000 human genes were knocked down with five or six short hairpin (sh)RNAs per gene in HeLa cells, and the change in sensitivity to montanine at two concentrations was quantified (Figure 5A and B). We found that sensitivity to montanine was significantly increased by the knockdown of four genes, including the EEF1A1 and GCN1L1 genes, which are related to protein synthesis (Figure 5C). The EEF1A1 gene encodes an isoform of the alpha subunit of elongation factor 1 (eEF1α), which recruits aminoacyl tRNA to the A-site of the ribosome during translation elongation. Knockdown (KD) of EEF2 mRNA, which encodes translation elongation factor 2, which is required for ribosomal translocation along mRNAs, also substantially sensitized cells to montanine (**5**) (Figure 5B). GCN1L1 encodes a human ortholog of yeast Gcn1 that senses ribosome stalling during translation (see Discussion). GCN1L1/GCN1 forms a complex with RWDD1 (also known as DFRP2, a human ortholog of yeast Gir2) and DRG2 (Rbg2 in yeast), the knockdown of which sensitizes cells to montanine (**5**) (Figure 5B) (25).

**Figure 5.**
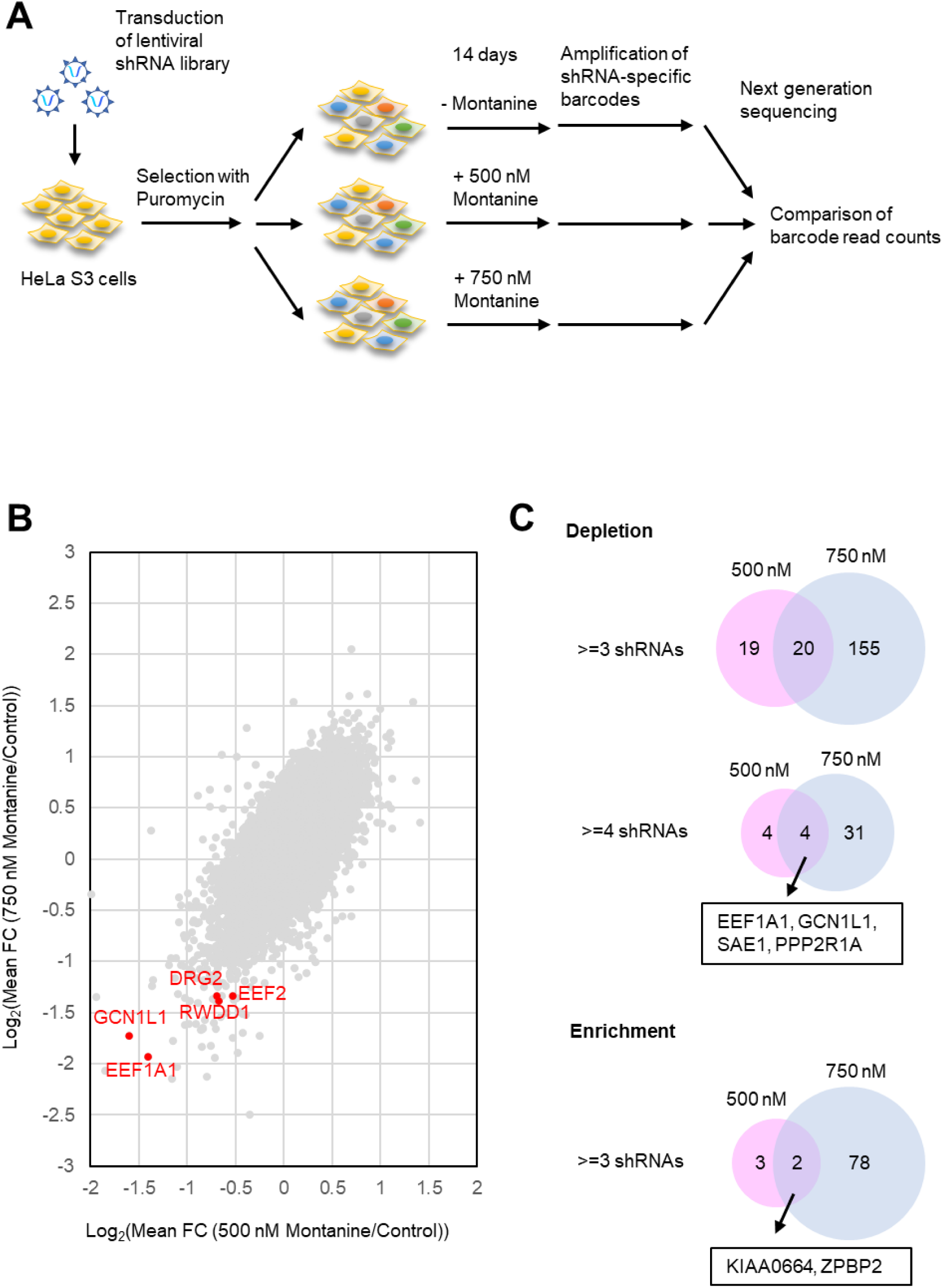
Chemical genomic analysis of montanine (**5**). (**A**) Schematic of the shRNA library screening. (**B**) The normalized read count of each shRNA from the montanine-treated samples was divided by that from the untreated samples (control) to obtain the fold change (FC, montanine-treated sample/control). The log_2_ (mean FC of shRNAs for each gene) values from the shRNA screens with 500 nM montanine and 750 nM montanine were plotted. (**C**) Comparison of candidate genes that determine the sensitivity to montanine between shRNA screens with 500 nM montanine and 750 nM montanine.

Our chemical genomic screening clearly suggested that the action of montanine involves the translational elongation step. Two previous independent functional genomics screens revealed that interference with the EEF1A1 and EEF2 genes significantly reduced DENV-2 replication in HeLa and HuH-7 cells (Tables S3 and S4) (26, 27). Similarly, KD of the RWDD1, DRG2, and GCN1L1 genes also impaired viral infectivity (Tables S3 and S4), suggesting them to be proviral host factors. Taken together, our analysis strongly suggested that the binding of montanine to ribosomes influence translation of viral mRNA governed by the above host factors.

## Discussion

In this study, we identified montanine (**5**) as a novel broad-spectrum antiviral effective against DENV, SARS-CoV-2, and Ebola pseudoviruses. Amaryllidaceae alkaloids (AAs) and their derivatives are well-established sources of antiviral compounds (13). Among the most extensively studied broad-spectrum antiviral AAs are lycorine (**1**) (28–31) and narciclasine (**2**) (32–34). Notably, montanine (**5**) demonstrated superior selectivity index (SI) values across all tested cases. Montanine-type AAs are known for their diverse biological activities, including antiproliferative and cytotoxic effects in various cancer and noncancer cell lines, as well as their antibacterial and antiparasitic properties (35). Interestingly, **5** exhibited antiarthritic activity in antigen-induced and collagen-induced arthritis models (36) and displayed behavioral activity related to inhibitory neurotransmission (37). Despite their wide range of bioactivities, the mechanism of action as well as the target molecules of montanine-type alkaloids have not been characterized. Therefore, the protein synthesis inhibition and ribosome-bound configuration revealed in the present study constitute the first report of the molecular target of montanine (**5**). The guanine-binding site in the PTC is also targeted by plant alkaloids, some mycotoxins (21), and the marine sponge alkaloid agelastatin A (24), where a small binding pocket is shared by seemingly structurally unrelated small molecules owing to the structural flexibility of ribonucleosides surrounding the binding pocket. However, our antiviral assays revealed that the methoxy group at C2 on the tetrasubstituted cyclohexene ring was critical for antiviral activity, as pancracine (**4**), a desmethyl derivative of **5,** exhibited weaker activity than did **5**. The α-stereochemistry of this methoxy group in **5** also plays a role since its epimer coccinine (**6**), with a β-stereochemistry at this functionality, results in significantly lower antiviral activity. The cryo-EM structure of the ribosome–montanine complex suggested that the oxygen atom and the methyl group of the methoxy group of **5** contributes to stabilizing the binding of the molecule via hydrogen bonding with U4450 and polar interactions with the phosphate moiety of G4451 in addition to several van der Waals interactions with surrounding nucleotides (Figure 4B). These interactions are important structural determinants of drug activity; thus, structural modification at this position would change the antiviral action of montanines. To optimize the functional moiety in the cyclohexene ring, the development of an efficient chemical synthesis scheme for **5** is highly desirable.

Inhibition of the host translation machinery to suppress viral protein synthesis might be expected to negatively impact host cell viability. However, eukaryotic cells have evolved antiviral responses that involve restricting global translation through the phosphorylation of the alpha subunit of eukaryotic initiation factor 2 (eIF2α). This mechanism activates the translation of a specific subset of mRNAs as part of the integrated stress response, ultimately triggering downstream antiviral defenses. Therefore, the inhibition of translation by small molecules may somehow mimic this antiviral response of the cell (38). We demonstrated that the knockdown of the translation factor eEF1A and stress-related GCN1 sensitized cells to montanine (**5**). Since eEF1A recruits tRNA to the A-site (39), its knockdown may synergize with PTC inhibitors, including montanine (**5**). Notably, ternatin-4 and plitidepsin, both of which target eEF1A, display inhibitory effects on SARS-CoV-2 (40–42).

GCN1 is known to be involved in the attenuation of translation upon stress, e.g., amino acid starvation, by the activation of another eIF2α kinase, GCN2, leading to the integrated stress response. Recent studies highlighted an additional role of GCN1 in ribosome quality control (RQC). Upon treatment with translational inhibitors or the presence of a stop codon readthrough, even under some normal conditions, slowing translation results in ribosome collision and stalling translation (25, 43, 44). In this process, GCN1 acts as a sensor for ribosome collision by forming a complex with colliding ribosomes (disomes) (45). Knockdown of GCN1 negatively affects RQC, which triggers the degradation of nascent peptides and the dissociation of ribosomal subunits from mRNA. It is possible that montanine-bound ribosomes might stall, and in the absence of proper RQC, aborted translation would result in sensitization of the cell to montanine treatment (Figure 6).

**Figure 6.**
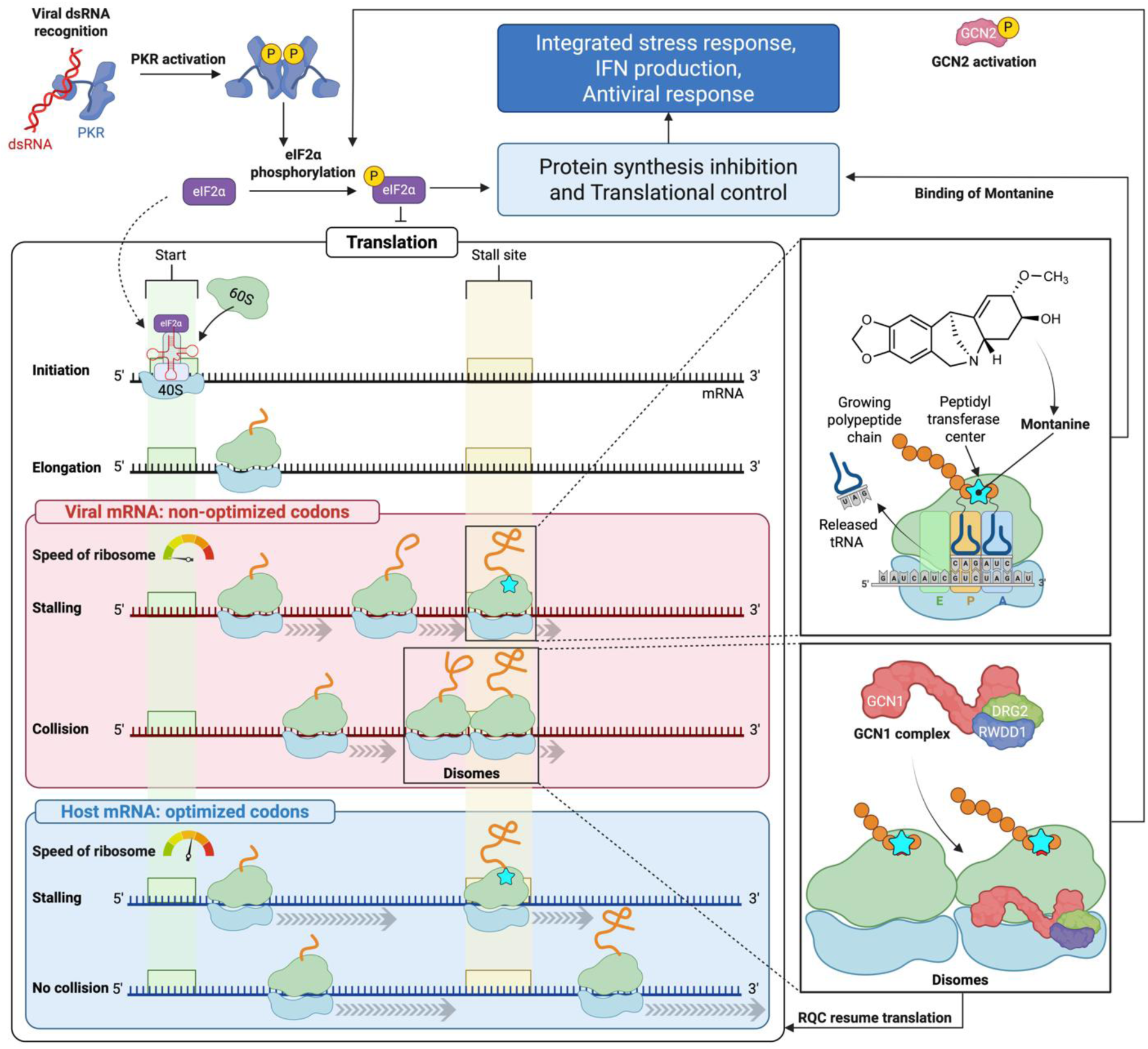
Proposed mechanism of montanine-mediated antiviral activity. Viral double-stranded RNA (dsRNA) activates PKR, leading to eIF2α phosphorylation and inhibition of translation. Concurrently, montanine binds to the ribosomal peptidyl transferase center (PTC), stalling elongation on viral mRNAs, which often contain non-optimal codons and exhibit slower ribosome movement than host mRNAs. This stalling induces ribosome collisions and the formation of disomes. Disomes are recognized by the GCN1 complex, which activates either the ribosome quality control (RQC) pathway to resolve the disomes or GCN2, further contributing to eIF2α phosphorylation. Together, PKR activation and montanine-induced ribosome stalling may converge to initiate the integrated stress response (ISR), promoting interferon (IFN) production and enhancing antiviral defense. In contrast, host mRNAs are translated more efficiently, reducing off-target effects and enabling selective inhibition of viral translation.

Notably, in recent functional genomic screenings to identify proviral factors for DENV infection, reanalysis of the data we mined EEF1A1 and EEF2 consistently . Additionally, we observed that suppressing the RWDD1, DRG2, and GCN1L1 genes also reduced viral infectivity suggesting that translational control mediated by the GCN1 complex influences viral replication. Our results indicate that montanine (**5**) targets the translational elongation step to inhibit viral replication while sparing host mRNA translation, as evidenced by its high selectivity index (SI) values.

Interestingly, a recent study revealed that the DENV and SARS-CoV-2 genomes preferentially use nonoptimal codons, possibly to modulate translation efficiency in host cells (46). This finding suggests that viral genomes are translated more slowly than host genes are translated, resulting in a greater dependency on translation elongation factors and RQC. Consequently, viral translation appears to be more sensitive to the effects of montelanine (Figure 6).

In summary, this study highlights translational control as a promising strategy to attenuate viral infectivity and a potential avenue for the development of small-molecule antivirals. Compared with the well-established antiviral agent lycorine, which also targets and inhibits ribosomes, monanine is an example of such a molecule and has superior activity. The distinct activity profiles of montanine (**5**) and other ribosome inhibitors suggest that the binding mode of small molecules can uniquely influence translational efficacy. Optimized compounds that target translation hold significant promise as effective antiviral agents with minimal host toxicity. In support of this concept, recent findings on plitidepsin revealed its multifaceted translational inhibition and broad-spectrum antiviral effects, further underscoring the potential of this approach. (47). These observations reinforce the concept that precise modulation of translational control could unlock new opportunities for developing novel BSAs. Advancing our understanding of structure–translation relationships will be essential in uncovering potent antiviral candidates, making this area of research a promising avenue for future breakthroughs.

## Materials and Methods

### Natural Products

The source, extraction, purification and chemical characterization of all the plant and fungal metabolites tested in this work are reported in the corresponding literature. Lycorine hydrochloride monohydrate (**1**) and narciclasine (**2**) were purchased from TCI and Caman Chemical, respectively. Tetraacetylnarciclasine (**3**) was obtained from **2** via acetylation with acetic anhydride in pyridine as previously reported (48, 49). Montanine (**5**) and coccinine (**6**) (15), cytochalasin B (**16**) (50, 51); cytochalasin A (**22**) (50); sphaeropsidin C (**17**) (52), cavoxin (**21**) (53, 54), higginsianin B (**7**) (55), sphaeropsidone (**13**) (56), pachybasin (**26)**, haagenolide (**8**) (57), inuloxins A and C, and α-costic acid (**24**, **9** and **28**) (58); fusicoccin (**18**) (59), ungeremine (**11**) (60), massarilactones D and H (**25** and **10**) (61), mellein (**20**) (62), seiridin (**27**) (63), cyclopaldic acid (**19**) (64), viridiol (**12**) (65), and *epi*-epoformin (**23**) (66) were isolated by A.E.

### Antiviral activity

High-content imaging analysis (HCA) was used to quantify DENV infection in a single-cell manner in HuH-7 cells. The cells were seeded at a density of 10,000 cells per well in PhenoPlate 96-well black plates and cultured overnight in Dulbecco’s modified Eagle’s medium (DMEM) supplemented with 10% heat-inactivated fetal bovine serum (FBS) and 1% penicillin/streptomycin (p/s) at 37°C with 5% CO_2_. The cells were infected with DENV-1 strain Hawaii (MOI of 5), DENV-2 strain 16681 (MOI of 1), DENV-3 strain H87 (MOI of 2), or DENV-4 strain H241 (MOI of 4) for 2 hours, and subsequently, the media were replaced with 100 µL of media containing test compounds at the desired concentrations. At 48 hours post infection (hpi), the cells were fixed with 4% paraformaldehyde, thoroughly washed with PBS (pH 7.4), and permeabilized with 0.1% Triton X-100 for 10 minutes at room temperature (RT). Blocking was carried out using Odyssey™ blocking buffer for 1 hour prior to immunostaining with the appropriate antibody. Viral RNAs in each cell line were visualized by immunostaining with the primary mouse monoclonal anti-dsRNA antibody, clone rJ2 (Merck) (1:1000) at 4°C overnight, and the secondary donkey anti-mouse IgG Alexa Fluor 647-conjugated antibody (Invitrogen) (1:1000) for 3 hours at RT. Nuclei were labeled with 300 nM DAPI (Invitrogen) for 30 minutes at RT. The fluorescence signals were captured by the Operetta CLS high-content system, and image analysis was conducted via the Columbus analysis system. DAPI signals were used for cell counting and calculation of cell viability. The percentage of infection was calculated by dividing the number of cells with positive viral RNA signals by the total number of cells.

Replication-competent recombinant VSVs were generated as described previously (67). VSV-G and VSV-ZGP were propagated in Vero E6 cells and stored at –80°C until use. The virus titers were determined via 50% tissue culture infectious dose (TCID_50_) assays. VSV-G and VSV-ZGP were inoculated into Vero E6 cells on 96-well plates (100 TCID_50_/well) together with the test compounds (1, 5, or 25 μg/mL), and the cells were incubated for 3 days. Cell viability was measured via the MTT (3-[4,5-dimethyl-2-thiazolyl]-2,5-diphenyl-2H-tetrazolium bromide) assay.

SARS-CoV-2 (68) was propagated in Vero E6-TMPRSS2 cells maintained in 2% FBS/DMEM at 37°C in 5% CO_2_. The infectious titers of SARS-CoV-2 were determined via TCID_50_ assays in Vero E6-TMPRSS2 cells. SARS-CoV-2 was inoculated into Vero E6-TMPRSS2 cells in 96-well plates (100 TCID_50_/well) together with the indicated concentrations of the test compounds, and the cells were incubated for 3 days. Cell viability was measured via the MTT assay.

Due to limited sample availability, tests was performed with n=1 (VSV-ZGP) or 2 (VSV-ZGP, SARS-CoV-2) but with three concentrations (25, 5, and 1 μg/mL).

### LysoTracker assay

HeLa cells were precultured overnight (3000 cells/well, glass bottom 96-well plate, Matsunami Grass), and an inhibitor (montanine 3 mM in DMSO, 0 mL) was added to the medium and incubated for 30 min. LysoTracker (Thermo Fisher, 000) was then added, and the mixture was further incubated for 30 min. The medium was removed, and the medium was changed to fresh DMEM. The cells were incubated under a fluorescence microscope (BZ 800, Keyence).

### Puromycin reporter assay

HeLa cells growing on a 96-well plate (50,000 cells/well, precultured overnight) were treated with inhibitors at the concentrations indicated above and incubated for 110 min. Puromycin (final concentration of 13.78 µM) was added to the culture and incubated for 10 min. The media was removed, the cells were washed with 1× PBST three times, and the cells were fixed in methanol. The methanol was removed, the cells were washed with 1×PBST three times, and the cells were blocked (Skim Milk, 60 min) and then washed with PBST three times. An anti-puromycin antibody (mouse, 1:5000 dilution, Merck, 12D10) was added to the cells, which were incubated for 1 h at room temperature, after which the cells were washed with PBST three times. The secondary HRP-conjugated anti-mouse IgG (1:5000 dilution; Cell Signaling Technology, #7076) was added to the cells, which were incubated for 1 h at room temperature, after which the cells were washed with PBST three times. One hundred microliters of TMB solution (Nakalai Tesque) was added, and the mixture was stirred for 30 s and allowed to react for 5 min. Then, 100 µl of reaction stopper solution (1 M sulfuric acid) was added, and the mixture was stirred for 30 s. The absorbance was measured in a microplate reader (450 nm).

### Chemical genomics shRNA screening

shRNA screening was performed as described previously (69, 70). Briefly, HeLa S3 cells cultured as described previously (70) were transduced with Decipher Human Module 1 (Cellecta, Mountain View, CA); one aliquot of transduced cells was untreated, and the other aliquots were treated with 500 nM and 750 nM montanine for 14 days and then collected. The same experiments were repeated with Module 2-infected cells and Module 3-infected cells. Genomic DNA extraction, PCR amplification of shRNA-specific barcodes with indexed primers, and barcode quantitation by next-generation sequencing on an Illumina HiSeq 2500 (Macrogen Japan) were performed as described previously (69).

Data processing was performed as follows. For each sample, the total number of reads was normalized to 20 million. shRNA-specific barcodes with fewer than 50 reads in the untreated samples were filtered out to exclude false positives. An additional 0.5 reads were added to the read count of each shRNA to ensure that all values were positive. The resulting read count of each shRNA from the montanine-treated samples was then divided by that from the untreated samples (control) to obtain a fold change (montanine-treated sample/control). Genes with <3 shRNAs and control genes in the libraries were excluded, and log_2_ (mean fold change of shRNAs for each gene) values were compared between shRNA screens with 500 nM montanine and 750 nM montanine. We selected candidate genes that determine the sensitivity to montanine with three or more shRNAs with thresholds of a fold change <0.5 (Depletion) and >2 (Enrichment).

### Cryo-EM sample vitrification and data collection

Montanine (30 μM final concentration) dissolved in DMSO was added to human 80S ribosomes (66 nM final concentration), purified from Expi293 cells (Thermo Fischer Scientific) as previously reported (71) via sucrose density gradient ultracentrifugation, and incubated at 37°C for 10 minutes. After incubation, 3 μL of this mixture was immediately applied to Quantifoil R1.2/1.3 Cu 200 mesh grids (Quantifoil) coated with handmade thin amorphous carbon that had been glow discharged at 10 mA for 10 sec with a sputter coater (JEOL JEC-3000FC). The samples were then vitrified via a Vitrobot Mark IV (Thermo Fisher) at 4°C and 100% humidity. Cryo-EM data were acquired via a CRYO ARM 300 II (JEOL) instrument operated at an acceleration voltage of 300 kV. A total of 5,525 movie micrographs were collected at 60,000x nominal magnification (corresponding to a physical pixel size of 0.788 Å) with a total dose of 40 electrons per Å^2^ via a K3 camera direct electron detector (Gatan) with the SerialEM program (72).

### Data processing

All data processing was performed via CryoSPARC2 version 4.1.2. (73). Briefly, patch motion correction and patch CTF estimation were performed on all the movies, and manually, the exposures were used to eliminate bad micrographs by thresholding and manual selection. Particle picking was performed on the selected micrographs via a template picker, which was extracted with a box size of 512x512 pixels, and then 2D classification was performed to select the particles. Ab initio reconstruction was performed on the purified particles, followed by homogeneous refinement to obtain the initial map. Focused 3D classification was performed for the low-density region of the 40S ribosomal subunit, and one class with distinct density was selected from the six classes obtained. After global CTF refinement and local CTF refinement, homogeneous refinement was performed again to obtain the final map of the human 80S ribosome in complex with montanine.

### Model building

The cryo-EM structure of the human 80S ribosome (74, 75) (pdb id: 4UG0 and 6z6m) was used for the initial model. The initial model of montmorillonite was prepared via ChemDraw (PerkinElmer Informatics, Inc.). The initial models were rigid-body fitted to the final map of the human 80S ribosome in complex with montanine via UCSF ChimeraX (76). Missing loops in low-density regions were manually deleted, and several rounds of manual fitting in COOT (77) and real-space refinement in Phenix (78) were performed to obtain the final structure. The statistics of the cryo-EM data collection, refinement, and validation are summarized in Table S4..

## Acknowledgments

This work was supported by JSPS KAKENHI Grant Numbers. 22H02430 to RS, 22H02252 to YT, and JP22H04922 (AdAMS to KM), the Japan Agency for Medical Research and Development (AMED, 20jm0210076h0002), and J-RAPID (20--201031622) to RS. SC was supported by the e-ASIA Joint Research Program from Thailand National Science and Technology Development Agency (NSTDA) and by the National Research Council of Thailand (NRCT; N33A670427, N35A660012). AT was supported by the Research Program on Emerging and Re-emerging Infectious Diseases from AMED (JP23fk0108624) and the Chemo-Sero-Therapeutic Research Institute (2023a018). YT and TY were supported by the Platform Project for Supporting Drug Discovery and Life Science Research (Basis for Supporting Innovative Drug Discovery and Life Science Research (BINDS)) from AMED under Grant Number JP23ama121038. Part of the work was supported by cloud funding READY FOR to RS.

## Supplementary Information

**Table S1.**
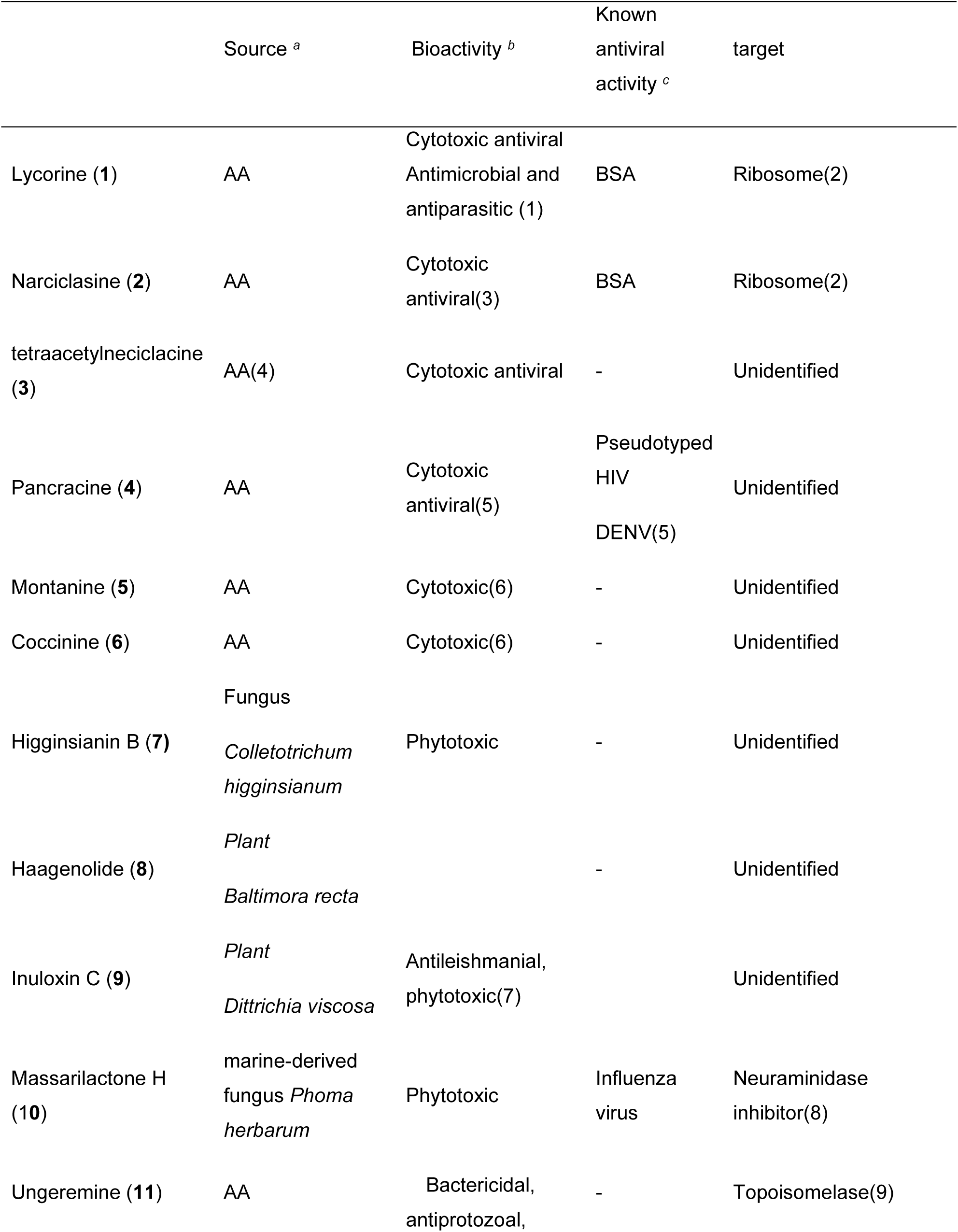
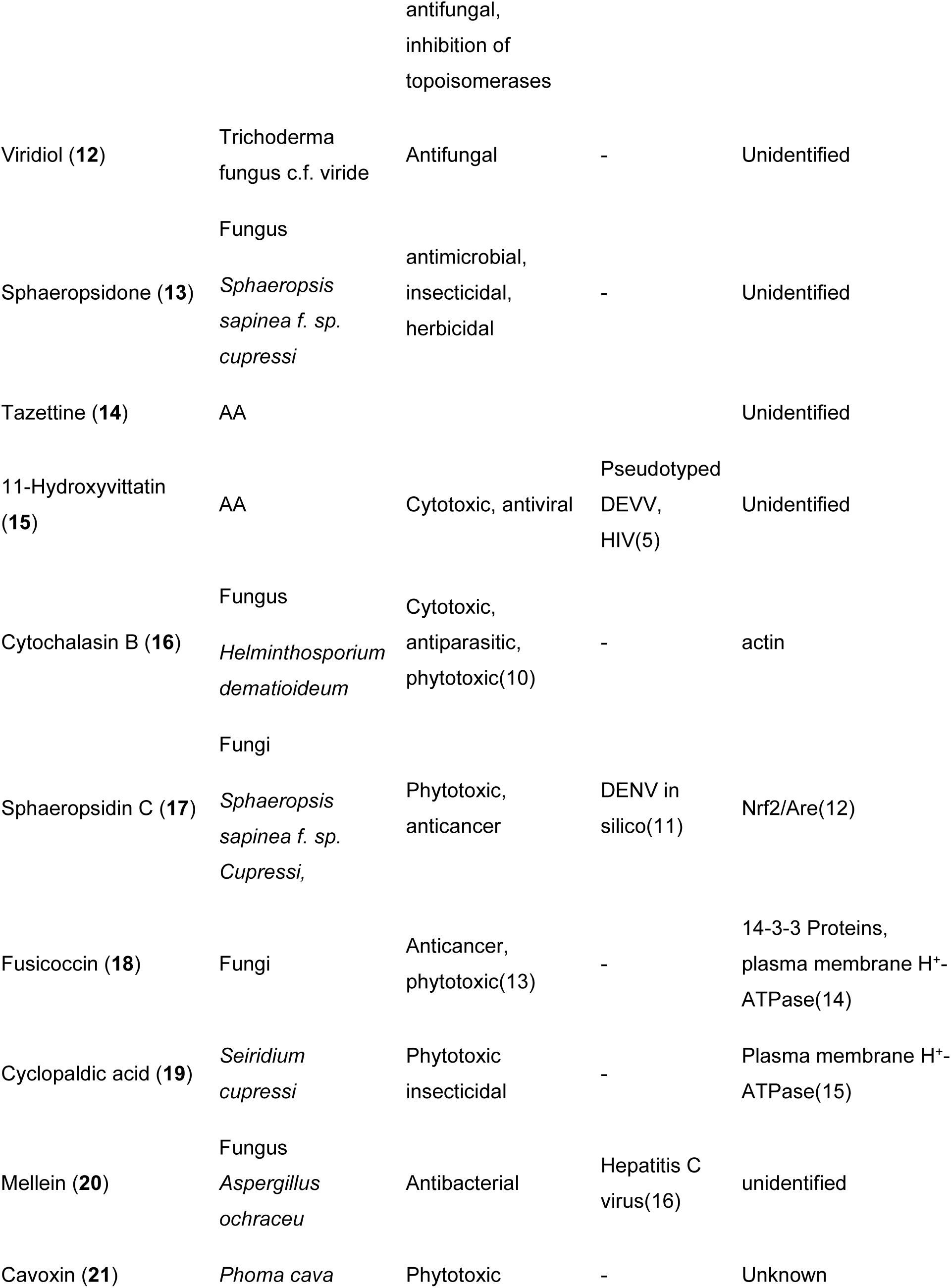
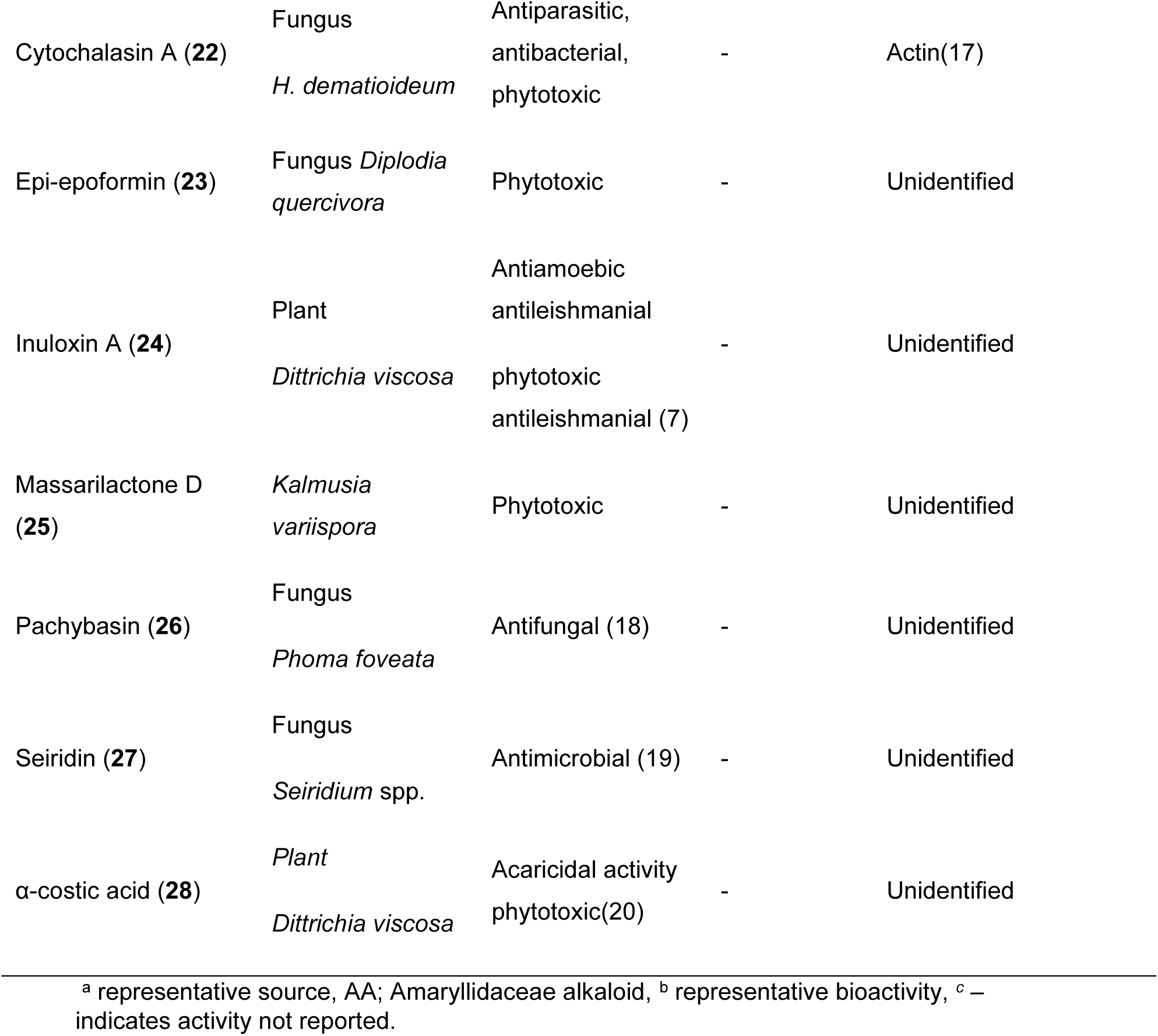
A library of plant and fungus-derived natural products.

### Targetted re-analyses of A genome-scale RNAi screen data (I)

A raw data set by Barrows, N.J. et al. (21) from genome-scale RNAi screen in HuH-7 cells using an siRNA library (22,909 mRNAs) identified host proteins essential for DENV-2 (New Guinea C strain) infection. EEF1A1, EEF2, RWDD1, DRG2, GCN1L1 were not discussed by Barrows et al, however, all above genes showed lower infection rates than the negative controls (GFP = 74.96% infection).

**Table S2.** A genome-scale RNAi screen in HuH-7 cells using an siRNA library (22,909 mRNAs) for pro-viral facots for DENV-2 (New Guinea C strain) infection.

| Gene | % Infection<br>(AB) | % Infection (CD) | Average | SD |
| --- | --- | --- | --- | --- |
| EEF1A1 | 15.29 | 43.87 | 29.58 | 20.21 |
| EEF2 | 28.54 | 55.11 | 41.83 | 18.79 |
| RWDD1 | 11.6 | 22.69 | 17.15 | 7.84 |
| DRG2 | 16.08 | 73.36 | 44.72 | 40.50 |
| GCN1L1 | 67.96 | 75.11 | 71.54 | 5.06 |

Each gene was targeted by four siRNAs in two pools (Set AB, Set CD). Negative control (AllStars: commercially available non-targeting siRNA control 86.22% infection), Negative control (Nonsilencing, 46.93 infection), Negative control (GFP, 74.96% infection), Positive control (ATP6V0C) (known proviral Factor, 6.95% infection)

### Targetted re-analyses of a genome-scale RNAi screen data (II)

Re analysis of data set by Savidis et al (22) for genomic RNAi screens in HeLa cells infected with DENV-2 (NGC strain) were conducted. As in re-analysis (I, Table S2), EEF1A1, EEF2, RWDD1, DRG2, and GCN1L1 genes exhibited lower average normalized infectivity than the proviral control ATP6V0B.

**Table S3.**
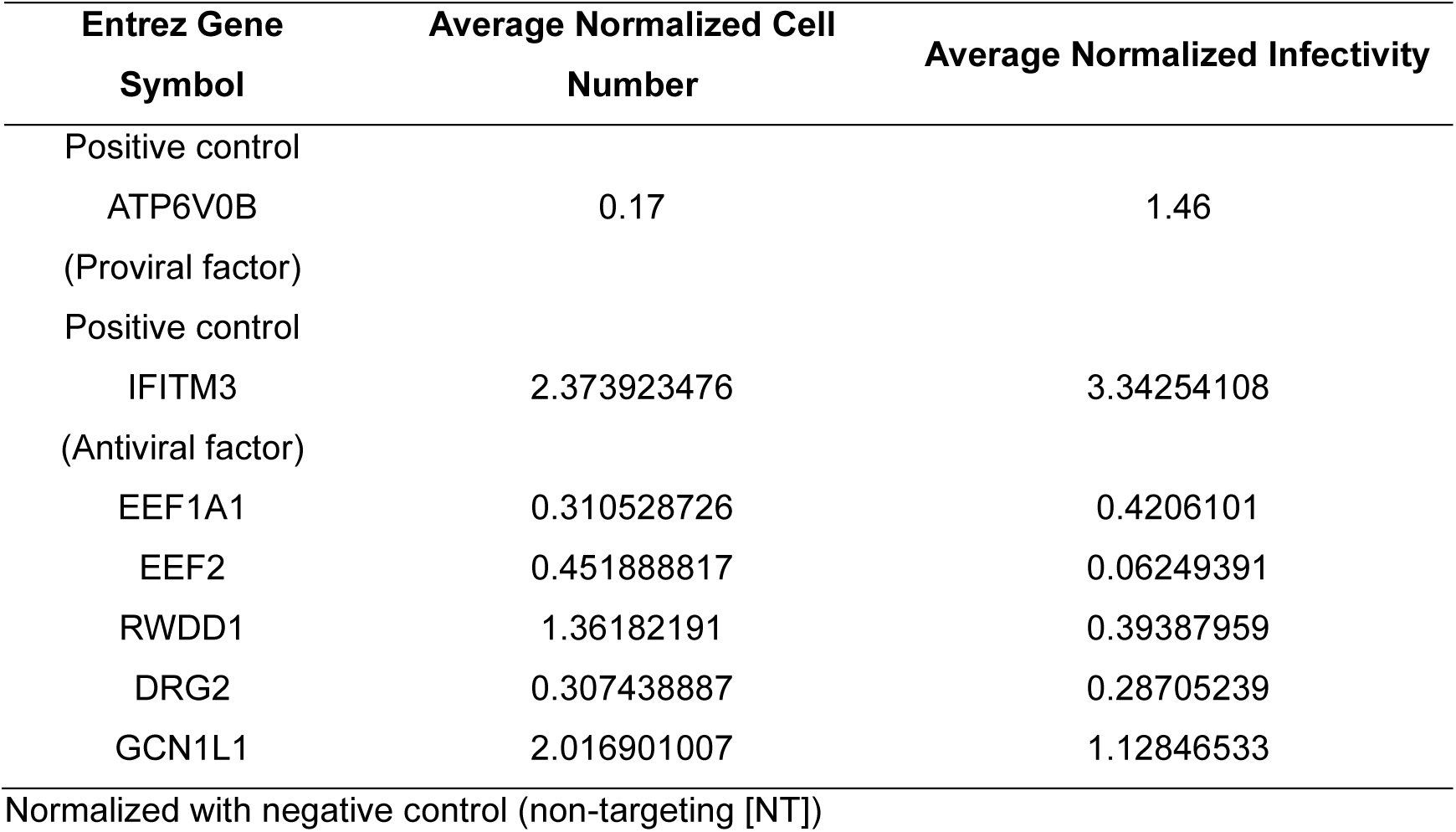
Genomic RNAi screens in HeLa cells infected with DENV-2 (NGC strain)

**Figure S1.**
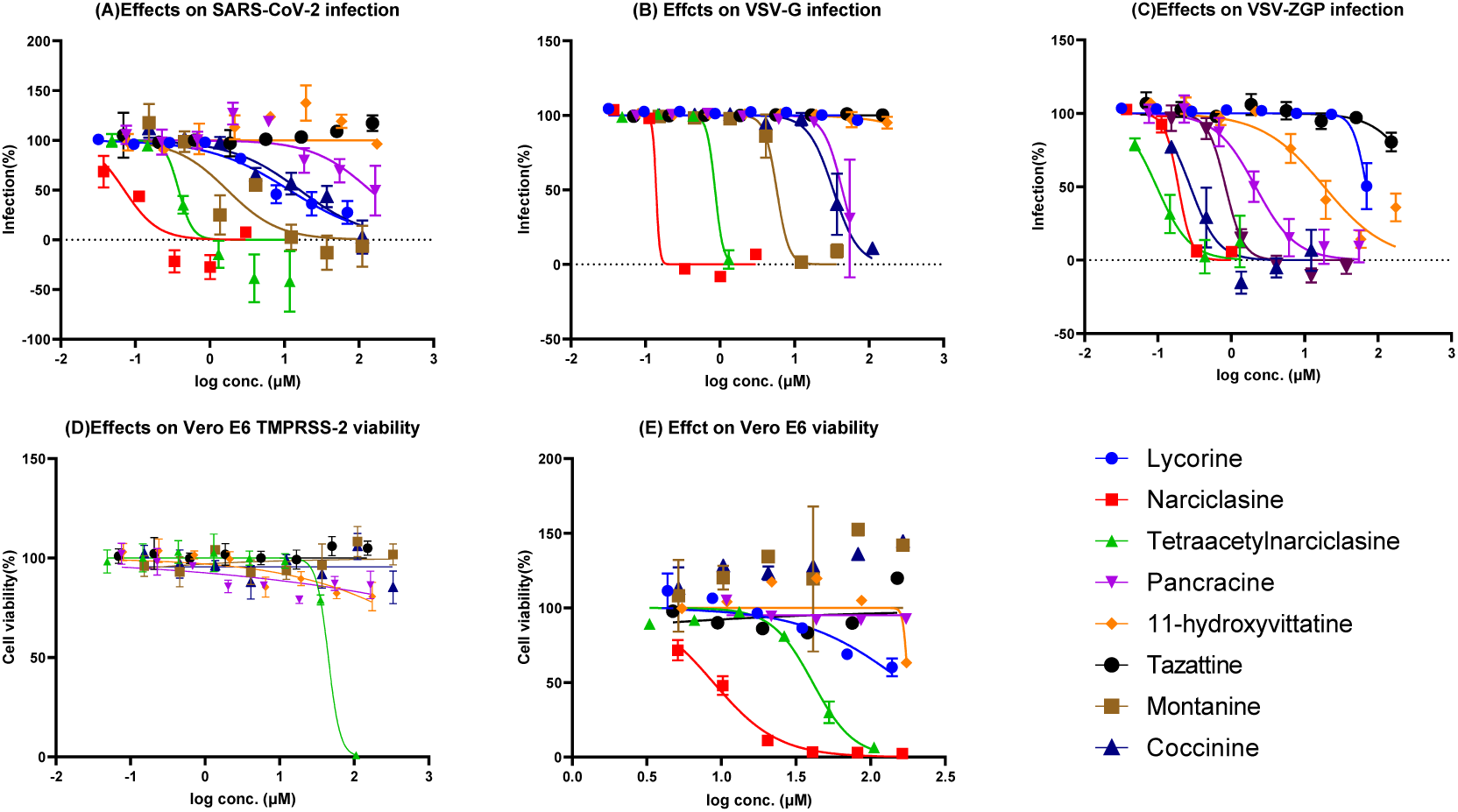
Concentration-activity relationship of Amaryllidaceae alkaloids on SARS-CoV-2, VSV and VSV-ZGP (C). Cytotoxocity aganist VeroE6TMPRSS-2 (D) and Vero-E6 cells (E) are shown.

**Figure S2.**
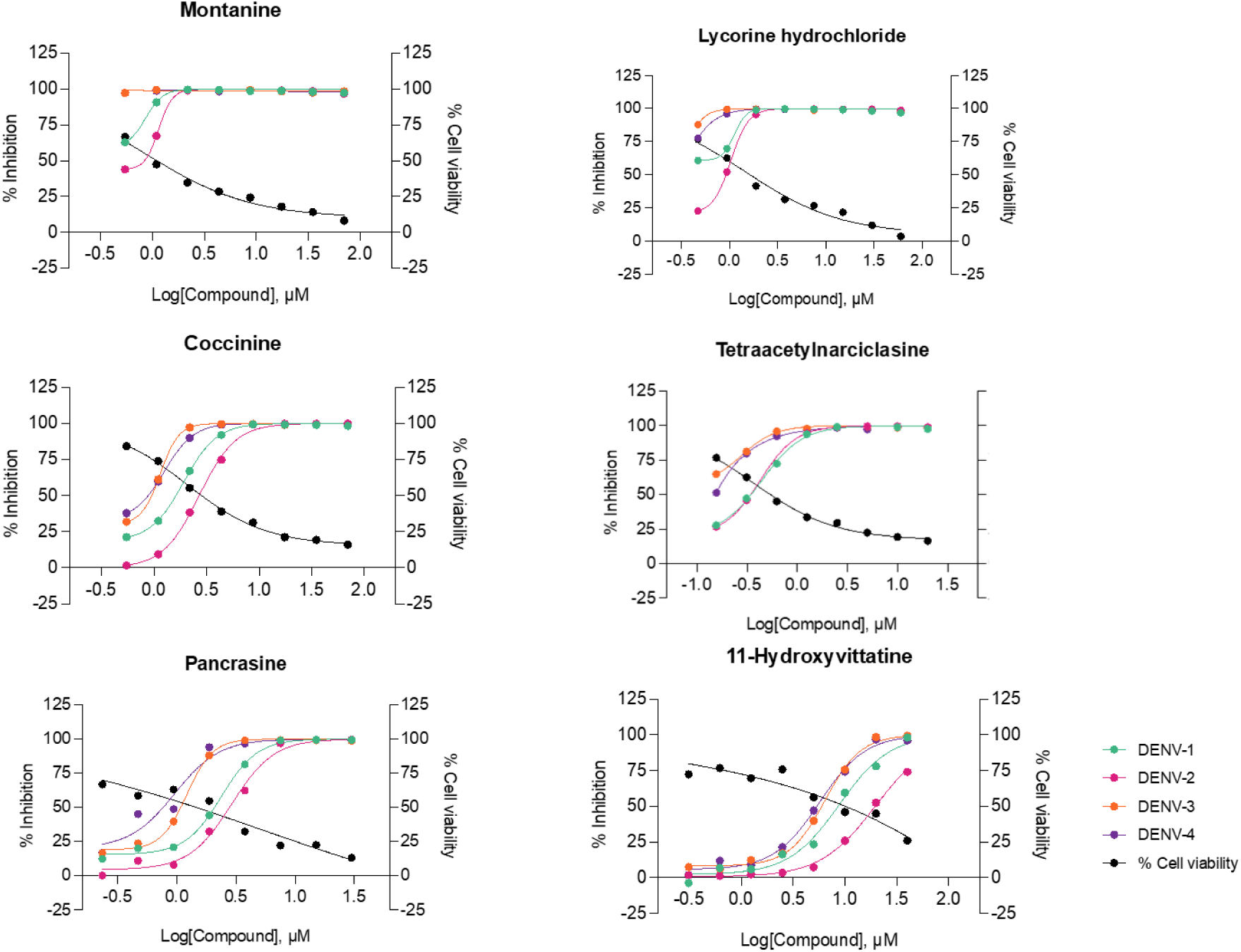
Concentration-activity relationship of Amaryllidaceae alkaloids on four serotypes of DENV. Cell viavilities were measured on HuH7 cells.

**Figure S3.**
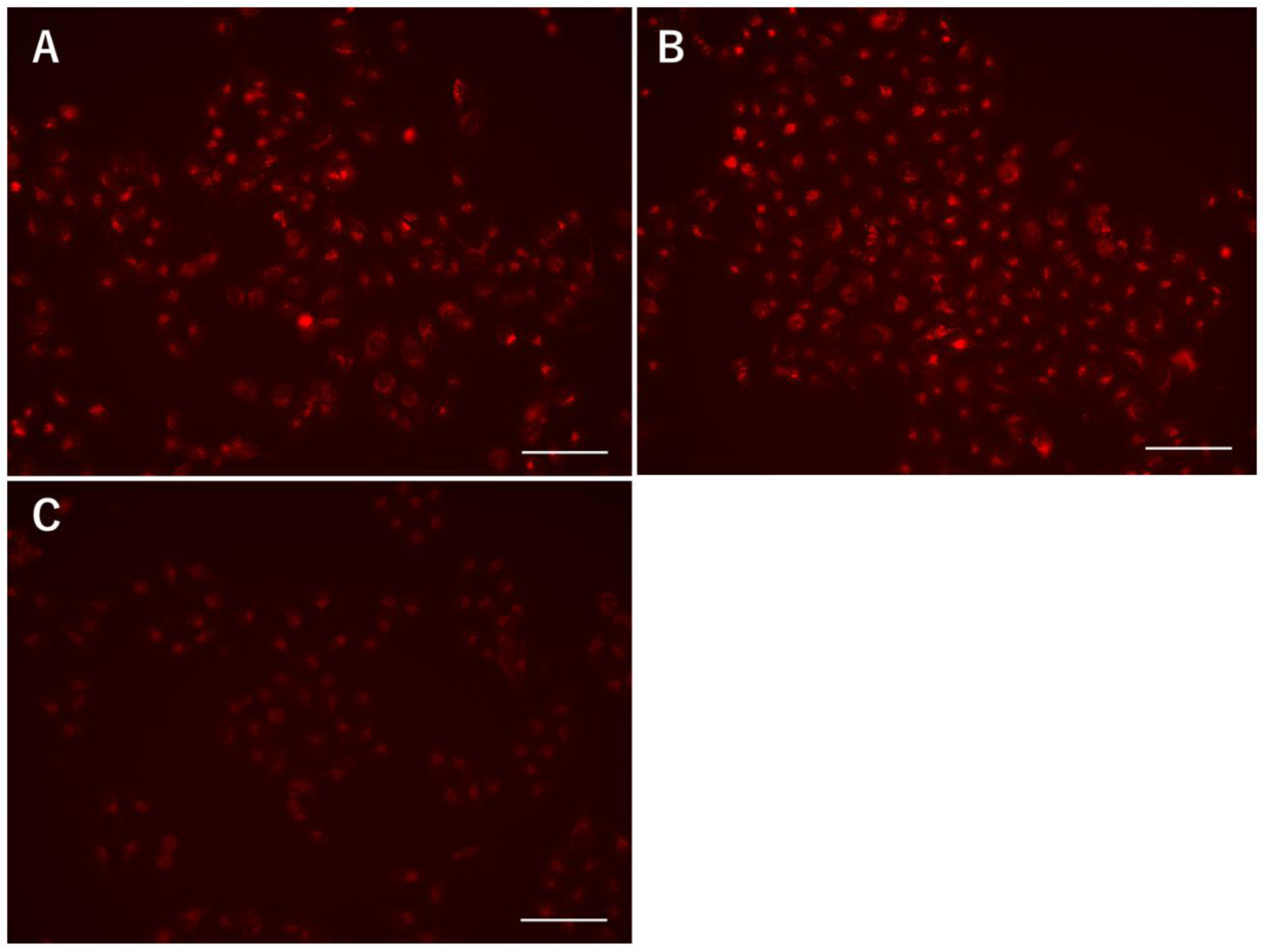
Fluorescent micrograph for HeLa cells treated with test compound followed by LysoTracker. (**A**) treated with montanine (3 μM), (**B**) negative control treated with DMSO. (**C**) treated with NH_4_Cl (10 mM).

**Figure S4.**
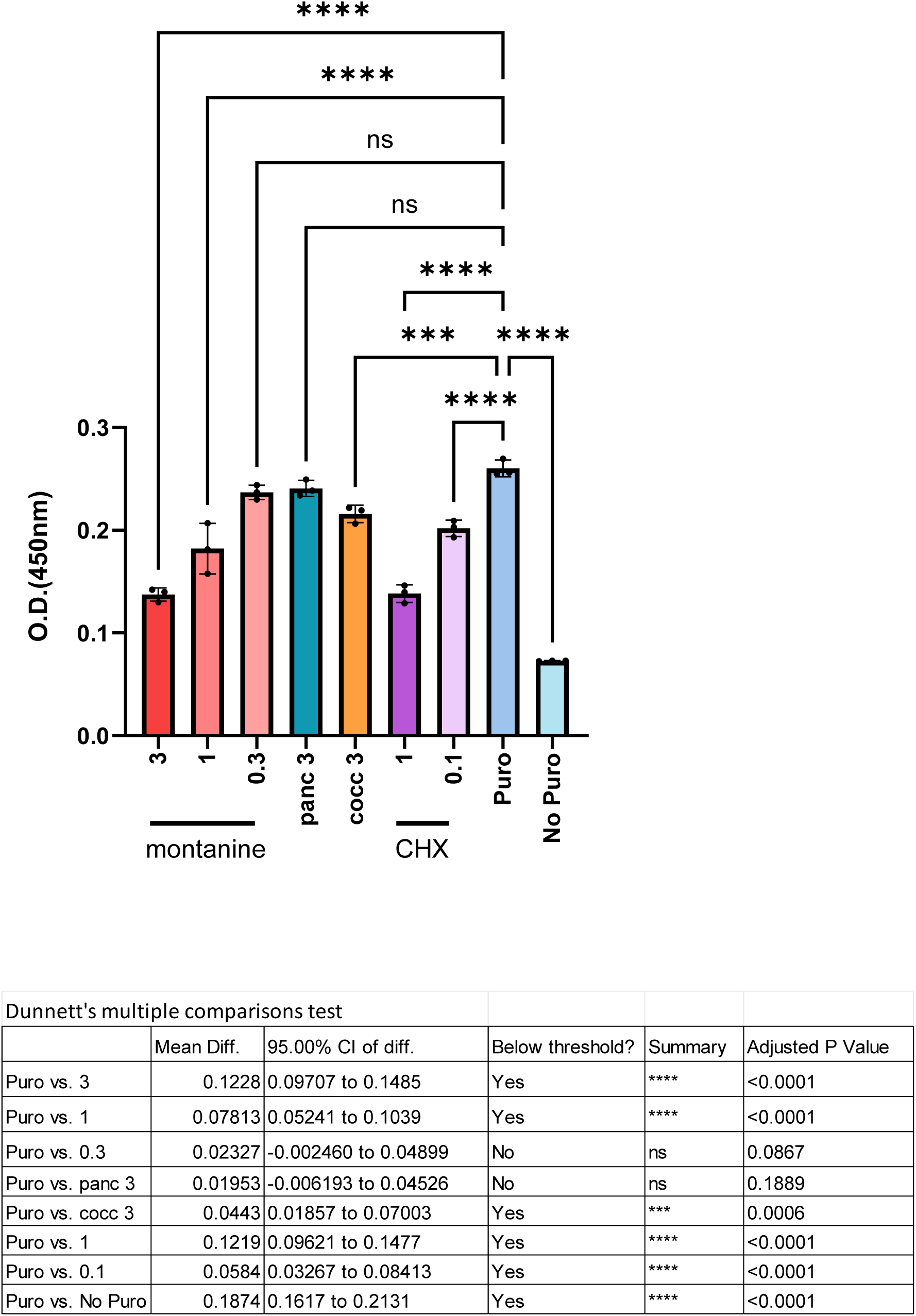
Statistical analysis using One-way ANOVA then Dunnett’s multiple **comparison.**

**Supplementary Figure S5.**
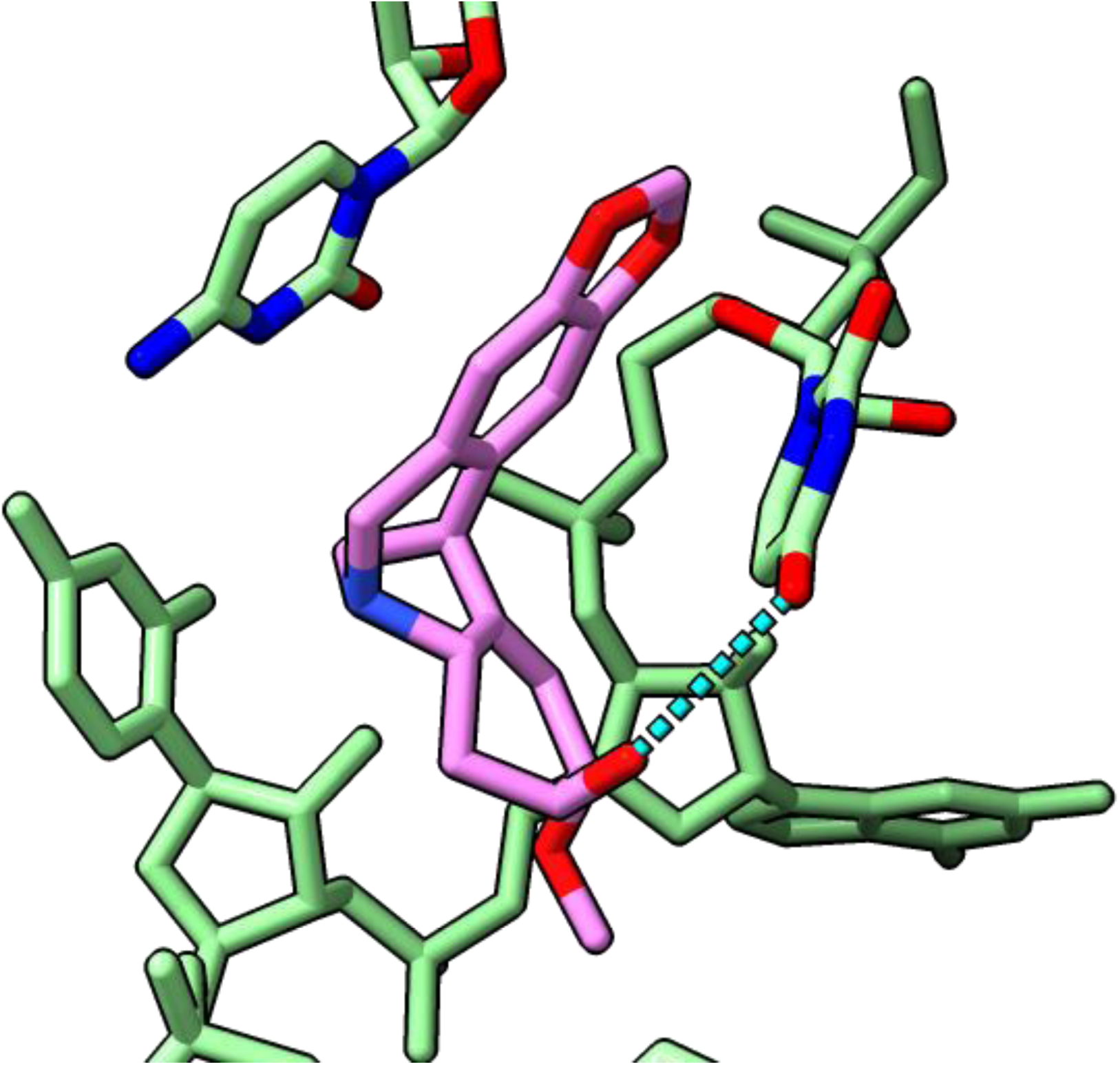
Sandwich-type π-π interaction by C4398 and U4452 and a hydrogen bond between montanine and U4552 Blue dotted line).

